# NeuroVelo: interpretable learning of temporal cellular dynamics from single-cell data

**DOI:** 10.1101/2023.11.17.567500

**Authors:** Idris Kouadri Boudjelthia, Salvatore Milite, Nour El Kazwini, Yuanhua Huang, Andrea Sottoriva, Guido Sanguinetti

## Abstract

Reconstructing temporal cellular dynamics from static single-cell transcriptomics remains a major challenge. Methods based on RNA velocity are useful, but interpreting their results to learn new biology remains difficult, and their predictive power is limited. Here we propose NeuroVelo, a method that couples learning of an optimal linear projection with non-linear Neural Ordinary Differential Equations. Unlike current methods, it uses dynamical systems theory to model biological processes over time, hence NeuroVelo can identify gene interactions that drive the observed temporal dynamics of gene expression. We benchmark NeuroVelo against several state-of-the-art methods using single-cell datasets, demonstrating that NeuroVelo simultaneously reconstructs correct cell-type transitions and identifies gene regulatory networks that drive cell fate directly from the data.

## 1 Introduction

Single-cell transcriptomic (scRNA-seq) technologies have transformed our understanding of cellular diversity and heterogeneity[1], yet the destructive nature of the measurement process poses fundamental limits to their ability to capture temporal biological dynamics. Inferring dynamic information from static snapshots of scRNA-seq has been a major focus of computational research in the last decade. While most early efforts used advanced machine learning techniques to order cells along a pseudo-time trajectory [2–5], more recently the concept of RNA-velocity [6] offered a more mechanistic, biophysically grounded approach to solve the problem.

RNA-velocity leverages the surprisingly abundant pre-mRNA reads present in many scRNA-seq data sets to parametrise a simple first order kinetic model of RNA production and splicing. The relative abundance of spliced and unspliced reads in a given cell can then be used to deduce whether a gene is being transcriptionally activated or repressed. This idea enables researchers to capture a shadow of cellular dynamics even in static data sets. Recent years have witnessed a flourishing of RNA velocity approaches [7–12], permitting a simultaneous inference of pseudo-time and RNA velocity, and coupling the velocity equations with non-linear transformations that can better model the complexity of the data. Nevertheless, both the biochemical foundations and the biological interpretation of RNA-velocity have been questioned [13, 14]. In particular, while RNA velocity enables an intuitive grasp of cellular dynamics at the level of individual cells, much less can be said at the level of the gene regulatory networks (GRNs) which underpin such dynamics, which in general can only be obtained by combining RNA-velocity analyses with post-processing with bespoke GRN inference methods.[15].

Here we propose NeuroVelo, a new approach which bridges the cellular and gene scales, simultaneously inferring temporal cellular dynamics and major gene interactions that drive such dynamics from scRNA-seq data. NeuroVelo is an approach grounded in dynamic system theory that combines ideas from Neural Ordinary Differential Equations (ODE) [16, 17] and RNA velocity in a physics-informed neural network architecture. We test extensively NeuroVelo on five scRNA-seq data sets and compare its performance against six recently published methods. On all data sets NeuroVelo achieved state of the art performance under a variety of metrics. Uniquely to our approach, we demonstrate how the interpretation of NeuroVelo results recovers highly accurate regulatory networks and biological processes underpinning the dynamics in a natural and mathematically consistent way.

## 2 Results

### 2.1 The NeuroVelo model

We first present a high-level overview of our method, focussing on its main features and innovative aspects. A detailed mathematical description of the model is provided in the Methods section. Figure 1(a) illustrates the basic concept of NeuroVelo. scRNA-seq data consisting of both spliced and unspliced reads are encoded to a low dimensional space through two dimensionality reduction channels: one is a non-linear 1D encoder learning a pseudo-time coordinate associated with each cell, while the second is a linear projection to an *effective phase space* for the system. The observed single-cell transcriptomes are modelled as discrete-time snapshots of a continuous time trajectory happening on this latent phase space.

**Fig. 1.**
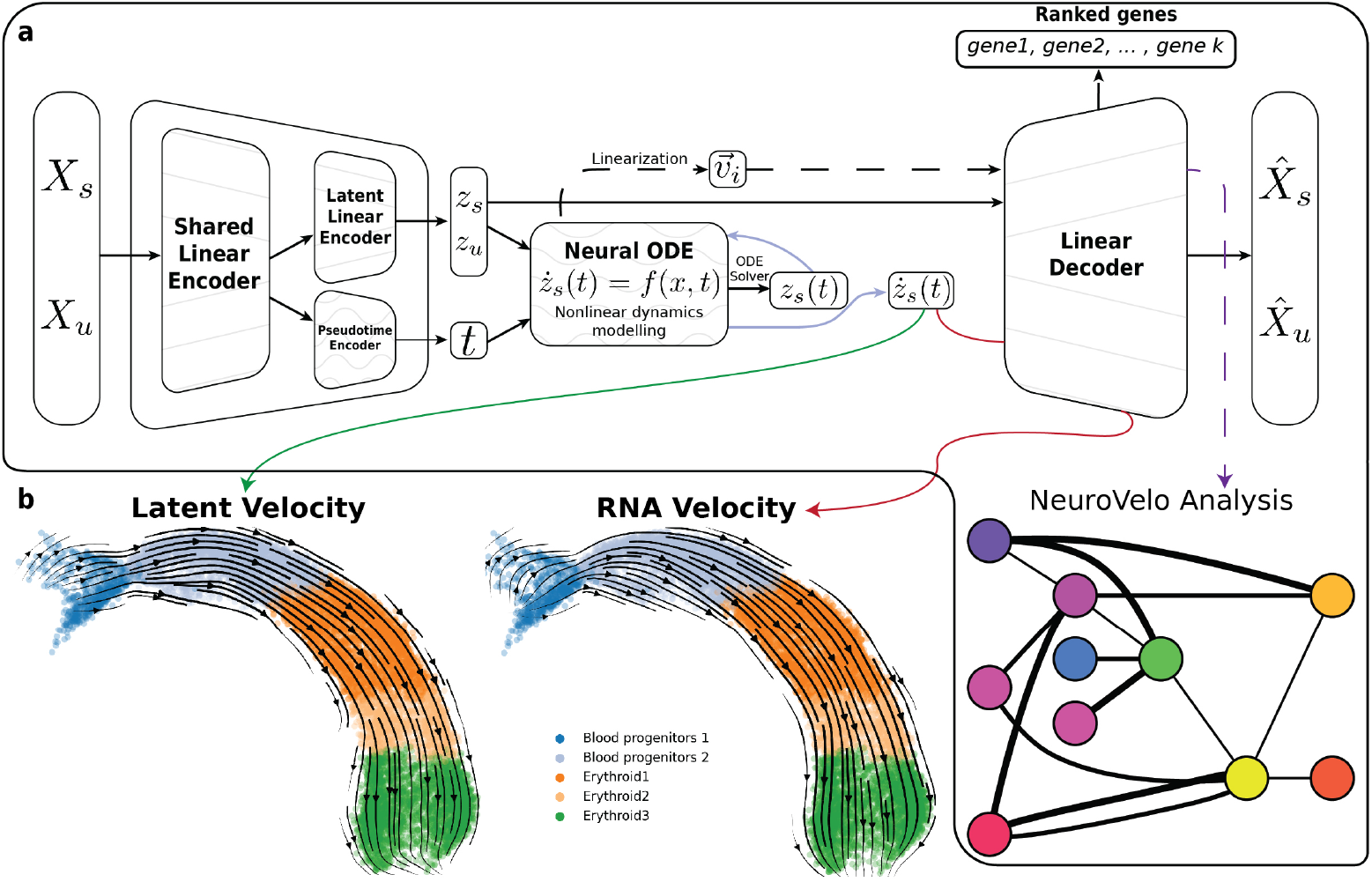
(a) Schematic representation of the neuroVelo model: two autoencoders, a linear one to define the phase space, and a non-linear 1-D one to define pseudotime, are coupled in latent space through a non-linear neural ODE. RNA-velocity constraints are added as an additional penalty in the loss function. (b) Example output of velocity field in the latent space and in the gene space on the mouse gastrulation dataset projected to the UMAP-based embedding

Cellular dynamics are defined through an autonomous system of differential equations where the rate of change of the state variables (right hand side of the ODE) is parametrised by a neural network (a neural ODE). The resulting dynamics can be extremely rich: since neural networks are universal function approximators, a neural ODE with a sufficiently large network may give rise to any first order dynamics. To ground biophysically these dynamics and avoid overfitting, we impose the RNA velocity principle as a further penalty in the loss function in the spirit of physics-informed neural networks. This introduces a trade-off in the fitting procedure: the parameters of the neural ODE and of the projections must strike a balance between providing an accurate interpolation of the observed transcriptomes, and maintain a balance of predicted spliced/ unspliced reads consistent with the RNA velocity principle. Notice that this constraint is applied locally to each cell, thus removing limiting assumptions on global transcriptional dynamics which are often a major problem for RNA velocity approaches. Because the effective phase space is defined via a linear projection, the RNA velocity constraint can be applied *exactly* in the reduced dimensional space.

The choice of defining the effective phase space as a linear subspace of gene space, while it may limit the expressivity of the model, brings additional significant advantages: it avoids confounding the non-linear dynamics in the effective phase space with the effects of a non-linear projection, leading to more robust results (see Supp. Fig. S1). Most importantly, a major benefit of the linearity of the phase space is that standard techniques for the analysis of low-dimensional non-linear dynamical systems can be applied to the trained model. The resulting insights, thanks to the linearity of the encoder/ decoder model, can readily be translated in terms of genes and regulatory interactions.

The main tool we use to interpret the regulatory interactions underlying the dynamics is the Jacobian matrix of the system. The Jacobian is obtained by taking first derivatives of the neural network with respect to the state variables, and it tells us how small changes in state variables affect their rates of change. The leading eigen-vectors -corresponding to the largest eigenvalues in magnitude-highlight the direction in phase space that contributes most to the system’s dynamics. In principle, both the Jacobian and its eigenvectors are time dependent. By decoding the Jacobian matrix in gene space, we can obtain estimates of how a gene affects the transcription rate of another gene at a particular time, hence providing a method to infer time-varying gene regulatory networks. Decoding the leading eigenvectors of the Jacobian into gene space provides ranked lists of important genes that can then be used for enrichment analyses and model validation.

### 2.2 Datasets used, competing methods and evaluation metrics

To test NeuroVelo, we utilize a total of five recent scRNA-seq data sets:

- a data set of erythroid cells differentiation [18] during mouse gastrulation, comprising of 9815 cells quantifying a median of 2791 non-zero expressed genes per cell;
- a human bone marrow hematopoiesis data set [5], widely used in RNA-velocity studies, which quantifies a median of 1190 non-zero expressed genes in 5780 individual cells;
- a pancreatic endocrinogenesis [19] data set measuring a median of 2447 non-zero expressed genes in 3696 cells;
- two smaller CRISPR-based lineage barcoded mouse cancer data sets [20], measuring gene expression in 754 and 1082 cells at a higher depth (3490 and 2352 median non-zero expressed genes respectively.

To contextualise the performance of NeuroVelo, we analysed each data set with a number of recently proposed methods to reconstruct cellular dynamics. Perhaps the most widely used RNA-velocity method is scVelo [7], which remains the main benchmark. Very recently, multiple methods extended the RNA-velocity framework in several directions: we choose here UniTVelo [9], which focusses on improving the pseudotime estimation, and latentVelo [21] and deepVelo [8], which couples RNA-velocity with non-linear dimensionality reduction. We also use scTour [17], which does not use RNA-velocity but directly fits a neural ODE to the data, as a splicing-independent comparison. On the lineage barcoded data, we use PhyloVelo [20], a specialised method which exploits barcode information to obtain robust estimates of pseudotime. As performance metrics, we use the cross-boundary direction (CBDir) and the in-cluster coherence (ICCoh), two widely used metrics which summarise the consistency of the inferred dynamics with known biological dynamics (in data sets where cell-type annotation is present). For all methods, we report the best performance across several random restarts, when a default initialisation is not provided. Table 1 provides a comprehensive assessment of all methods on all data sets with respect to the inferred RNA-velocity fields.

**Table 1.**
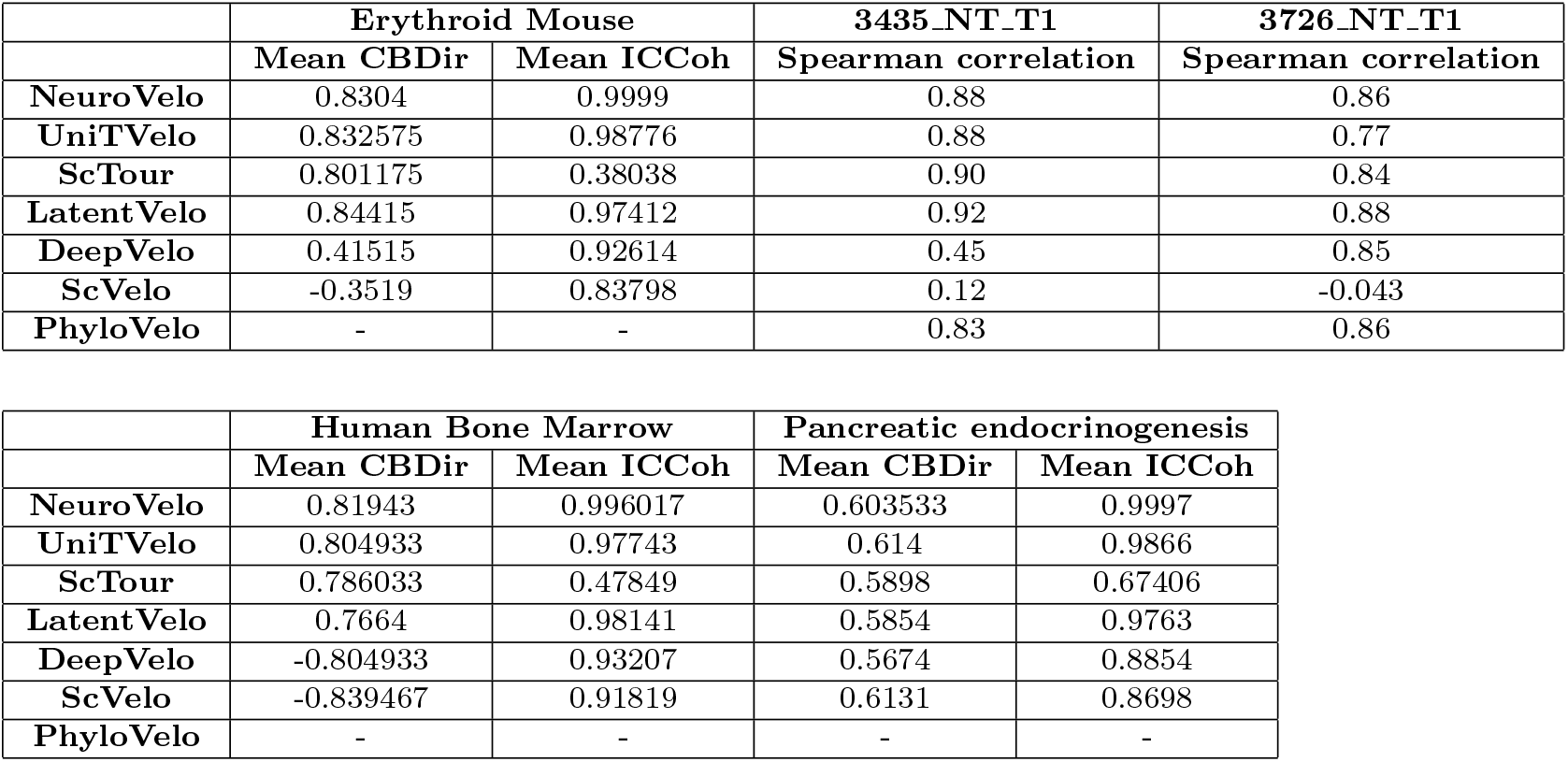
Metrics table for different methods and datasets. For method that allows different random seeds (NeuroVelo, ScTour and LatentVelo) the table shpws the best performance of running 10 seeds. More on stability of velocity is shown in figure S1

To benchmark the ability of NeuroVelo to reconstruct GRNs, we compare its performance to GENIE3 [22] and GRNboost [23], two machine learning-based methods which were highlighted as state of the art in a recent review [24]. Assessing the performance of GRN methods on real data is notoriously difficult, due to the lack of gold standards (and indeed the difficulty in associating precise molecular semantics to the interactions summarised by a network edge). To overcome this hurdle, we assembled a data base of ChIP-seq tracks from ChIP-Atlas [25], a number of transcription factors in a number of cell types which are relevant in gastrulation and hematopoiesis, the two most studied processes we consider. A list of all data sets used with accession codes is provided in supplementary table ST 1 and ST 2. This enables us to obtain a ground truth network for each cell type, where the various methods can be assessed using standard metrics such as area under the precision-recall (PR) curve.

### 2.3 Reconstructing dynamical mechanisms from scRNA-seq data

We start by assessing NeuroVelo performance on the three standard (non-barcoded) scRNA-seq, focussing on the ability to reconstruct lineage differentiation and GRNs. An analysis of the erythroid lineage in mouse gastrulation is shown in Figure 2.

**Fig. 2.**
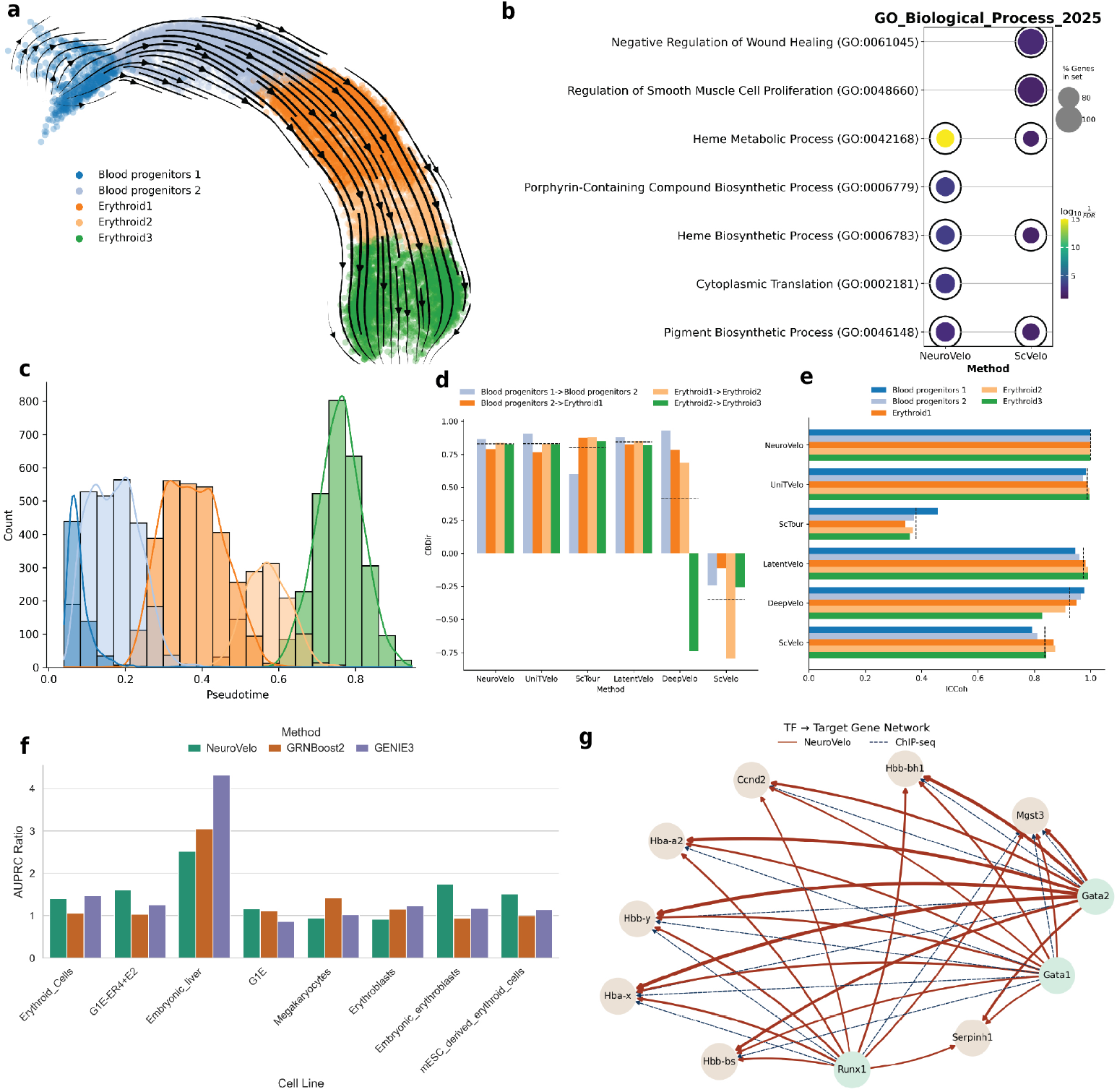
(a) Velocity field in gene space on the mouse gastrulation dataset projected to the UMAP-based embedding. (b) Comparison on gene set enrichment analysis of the gene influencing the dynamics. (c) Histogram of Inferred pseudotime. (d) Comparison on the cross boundary direction. (e) Comparison on the inter-cluster coherence. (f) Comparison on AUPRC ratio for the gene regulatory network inference. (g) Gene regulatory network inferred by NeuroVelo along with connections from Chip-seq data.

NeuroVelo effectively captures the single-branched nature of the process, providing the right cell-type transitions of the dynamics both in latent space and in gene space (Figure 2a), and a very good sequential ordering of the different cell types (Figure 2c). The quality of the inferred dynamics is further supported by Figure 2(d-e), which plots histograms of the evaluation metrics for each cell type, clearly showing NeuroVelo to provide state-of-the art performance on every cell type.

Figure 2f-g instead concern the interpretation of the decoded dynamics. The barplots in Figure 2f show the normalised area under PR curve for NeuroVelo, Genie3 and GRNboost for eight cell types which are associated with gastrulation (the normalisation is such that the random predictor would return a score of one). NeuroVelo returns the best performance in half of the cases, with a particularly marked improvement for two of the most relevant cell lines, embryonic erythroblasts and mESC derived erythroid cells. It should be noticed that NeuroVelo is agnostic with respect to which genes are transcription factors, while both Genie3 and GRNboost require as input the list of transcription factor genes, thus significantly narrowing the search space for their models. Figure 2b instead shows an enrichment analysis for NeuroVelo (using the ranked list of genes provided by the eigenvectors of the Jacobian) and for scVelo, using the list of driver genes in its dynamical model setting. The two analyses broadly overlap (see also Suppl. Fig. S3 for a comparison of the two gene rankings), although neuroVelo returns larger gene sets with a slightly higher significance level.

A similar picture emerges from the analysis of the human bone marrow data set in Figure 3. NeuroVelo provides the right cell-type transitions and a correct pseudotemporal ordering (Figure 3(a,c)), mirrored in state-of-the-art performance under both metrics (Figure 3 d-e). An enrichment analysis returns several highly enriched pathways related to immune cell proliferation and differentiation (Figure 3b), while in this case the gene ontology analysis was not able to identify meaningful biological signals from the scVelo driver genes set.

**Fig. 3.**
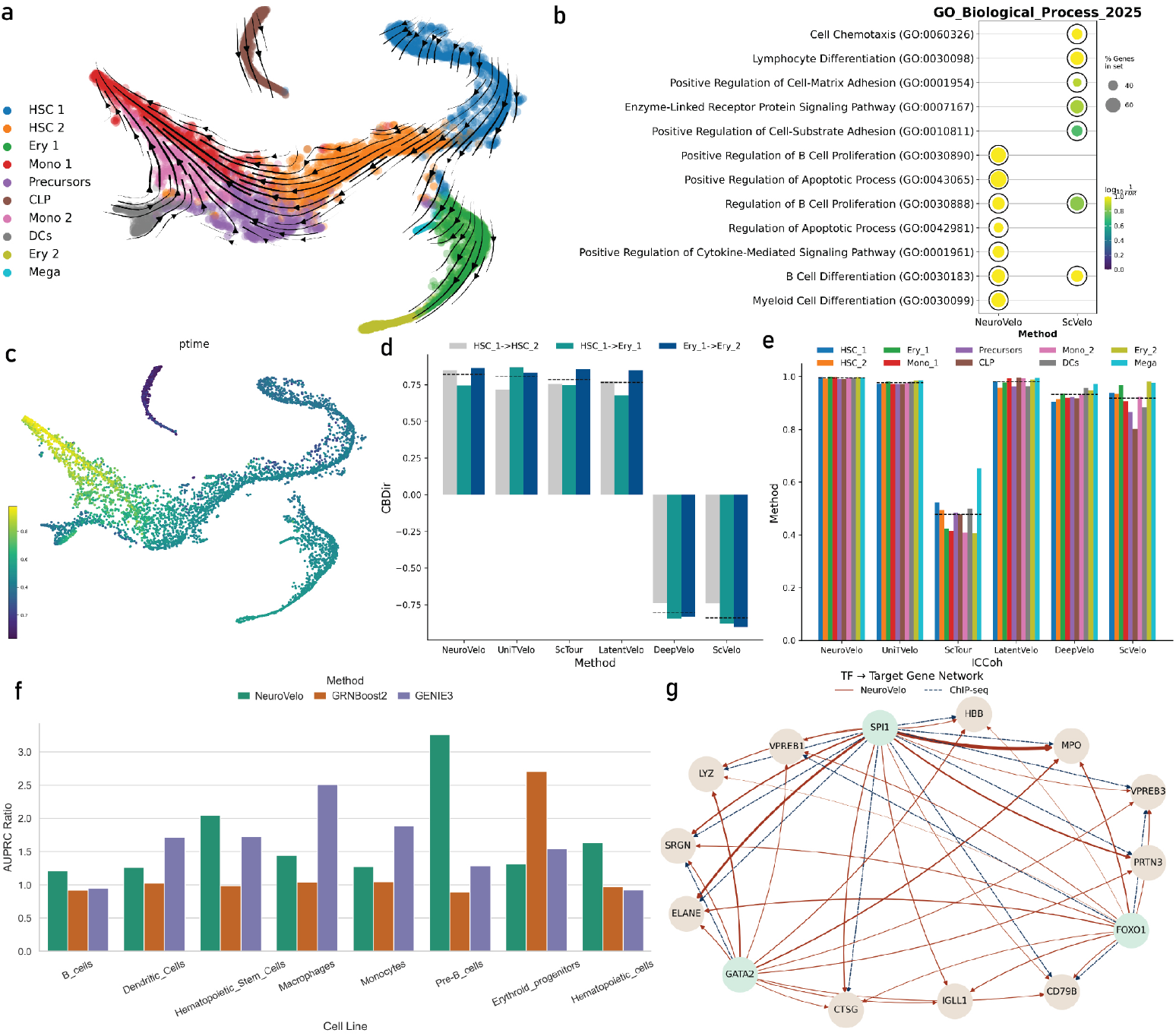
(a) UMAP of NeuroVelo velocity field on the human bone marrow dataset, as well as (c) NeuroVelo pseudotime. (b) Comparison on gene set enrichment analysis of the gene influencing the dynamics. (d) Comparison on the cross boundary direction. (e) Comparison on the inter-cluster coherence. (f) Comparison on AUPRC ratio for the gene regulatory network inference. (g) Gene regulatory network inferred by NeuroVelo along with connections from Chip-seq data.

Figure 3f again shows the agreement of the inferred networks with the ChIP-seq derived gold-standard in a number of relevant cell types. NeuroVelo achieves best performance in four out of eight cell types, including the most relevant cell types (hematopoietic cells and pre-B cells). Interestingly, in this well annotated data set we can also linearise about different points in latent space, producing cell-type specific networks which highlight the varying temporal dynamics in the process (see Supplementary Movies).

As a third example of neuroVelo analysis on standard scRNA-seq, we examined the pancreatic endocrinogenesis data set. Results are shown in Figure 4; once again, neuroVelo provides an excellent visualisation (Figure 4a), and performs best or among the best for all cell types under the metrics considered (Figure 4c-d). An enrichment analysis (Figure 4 b,e) highlights highly relevant pathways connected with pancreas function with very high normalised enrichment score. An analysis of the interactions predicted again highlights some interactions already detailed in the literature (Figure 4 f-g), however it is noticeable that in this less well studied scenario fewer interactions can be derived from the literature, underscoring the potential for data-driven approaches like neuroVelo to uncover novel biological hypotheses.

**Fig. 4.**
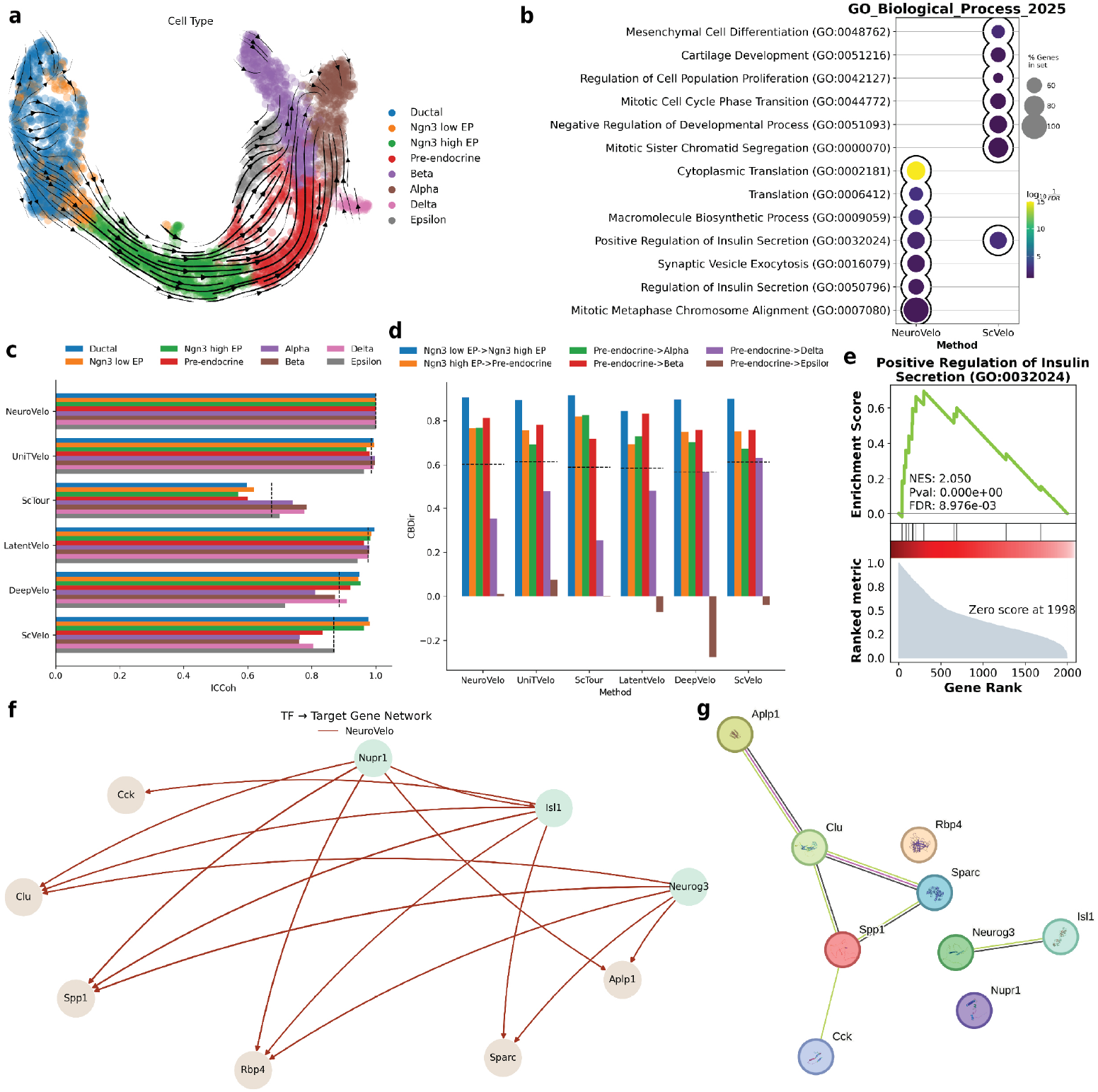
(a) UMAP of NeuroVelo velocity field on pancreatic endocrinogenesis dataset.(b) Comparison on gene set enrichment analysis of the gene influencing the dynamics. (c) Comparison on the in-cluster coherence. (d) Comparison on the cross boundary direction. (e) Gene set enrichment analysis of insulin secretion pathway. (f) Gene regulatory network infered by NeuroVelo and (g) gene network structured using string-db (right)

Taken together, these results demonstrate that neuroVelo achieves state of the art or is competitive with the best RNA-velocity methods on a variety of data sets. Additionally, neuroVelo is able to predict (possibly time-varying) interaction mechanisms which can explain the observed transcriptome dynamics, and which frequently recapitulate experimentally observed interactions in well characterised systems.

### 2.4 Inference of fitness in lineage barcoded data

As a second set of benchmark data, we consider two lineage barcoded data sets which were recently published by [26]. This technology combines scRNA-seq with barcoding to determine simultaneously the fitness of a cell population and its transcriptomic profile. [20] developed also a specialised “velocity” method called PhyloVelo, which does not use splicing information but relies on the barcoding to infer cellular dynamics. Results of the analyses on these two data sets are shown in Figure 5, which again show a very good visualisation on both data sets (panels a-b and d-e). To evaluate quantitatively the results, we focus on correlating the inferred pseudotime with the fitness signature, which is a specific gene module whose expression is associated with a cell’s proliferating fitness and it is estimated from the phylogenetic tree (see [26] for details on how fitness signatures are computed). Panel c shows the scatterplot of fitness values versus inferred pseudotime for one of the data sets, while panel f shows a comparison of all methods (including PhyloVelo) in terms of Spearman correlation with fitness. This shows that most methods perform extremely well (in fact, comparably to PhyloVelo, which explicitly uses the barcoding information), with the exception of scVelo and DeepVelo, this latter method only on one of the data sets.

**Fig. 5.**
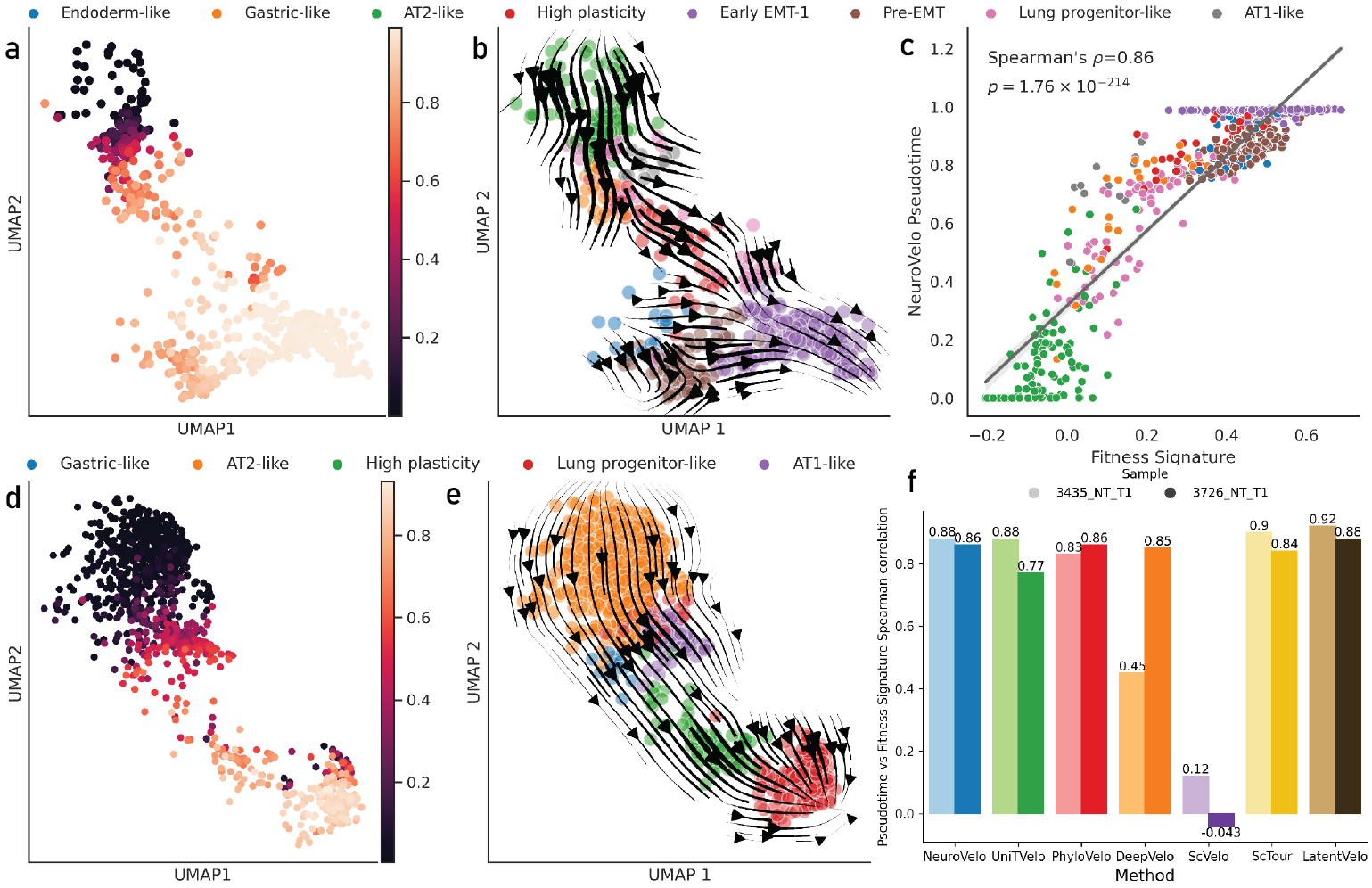
(a) UMAP of NeuroVelo infered pseudotime and (b) NeuroVelo velocity field on the sample **3726 NT T1** of the mouse cancer data set, as well as (c) scatterplot of the neuroVelo pseudotime against the fitness signature (inferred from barcoding information). (d) UMAP of NeuroVelo infered pseudotime and (e) NeuroVelo velocity field on the sample **3435 NT T1** of the mouse cancer data set. (f) Comparison on pseudotime and fitness signature spearman correlation for 2 different samples.

This analysis demonstrates that utilising splicing information can provide an effective way to reconstruct cellular ordering which validates against barcoding information. Once again, neuroVelo emerges as one of the strongest performers also on this task.

## 3 Discussion

NeuroVelo innovates over other RNA-velocity methods by bringing a new dimension of functional interpretability: while standard methods focus on defining a cell-level velocity field, depicting the evolution of individual cells, neuroVelo provides a data-driven mechanistic model which can be interpreted in terms of (time-varying) regulatory networks. Results on all data set demonstrated not only that neuroVelo is among the best performing tools in terms of standard RNA-velocity metrics, but also that the mechanisms it captures are biologically meaningful and extend known interactions from the literature. Importantly, a quantitative assessment of the reconstructed GRNs against experimental ChIP-seq data showed NeuroVelo to perform competitively with bespoke GRN inference methods which were supplied the list of transcription factor genes, a notable advantage which neuroVelo evidently could reconstruct independently from data.

Methodologically, neuroVelo attempts to strike a balance between complexity and interpretability. It does so by limiting its non-linearity to the dynamical system in the low-dimensional latent space, which enables both a very direct interpretation in terms of genes and the imposition of the RNA-velocity constraint as physics-informed loss term. An additional benefit of the relative simplicity of neuroVelo is its robustness.

Defining a semi-mechanistic model for single-cell dynamics opens a number of exciting opportunities to expand the scope of neuroVelo. First of all, principled ways to incorporate known mechanisms might be devised using ideas from Bayesian statistics. Secondly, other data modalities, such as chromatin accessibility, could be incorporated when available, thus providing a mechanistic extension to current methods for multiomics analysis [27]-[28, 29]. Finally, different types of underlying dynamics might be considered, for example to accomodate mutational events which might lead to non-smooth dynamics, of to incorporate spatial correlation in spatial transcriptomic data.

## Methods

### RNA velocity

ScRNA-seq captures a static snapshot of gene expression at a single time point, hence modeling cellular dynamics from single-cell RNA-seq data is not straightforward. La Manno et al. [6] leverage the presence of unspliced pre-mRNA to model transcriptional dynamics from single cell data. The coexistence of unspliced and spliced transcripts enables the inference of RNA velocity by modeling the underlying splicing kinetics. A simple model was assumed by [6] describing the evolution of unspliced and spliced mRNA molecules over time given by the system of equations 1

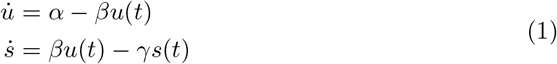

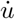 and 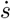 are time derivatives of the abundance of unspliced and spliced RNA (also known as RNA velocity), respectively. *α* is the transcription rate, β is the splicing rate, and γ is the degradation rate of the spliced RNA. By estimating parameter β and γ, one can make RNA velocity estimate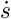, which is a vector in gene space that gives the infinitesimal change in abundance of spliced RNA amount (hence the analogy with the concept of velocity from classical mechanics).

### Model architecture

NeuroVelo is a neural network architecture that consists of a linear auto-encoder and neural ODEs architecture in the latent representation of the data. The main idea of NeuroVelo is to construct cell dynamics from scRNA-seq data in the latent space.

### Encoder/ decoder architecture

We used spliced and unspliced reads of scRNA-seq data with *g* number of genes, and embedded them using the same 2 layers linear encoder function *E* : ℝ^*g*^ → ℝ^*l*^, into a latent space ***z*** = (***z***_*s*_, ***z***_*u*_) of dimension *l*.

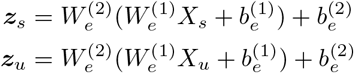

This step is mainly for dimensionality reduction and feature extraction of the highly dimensional gene space to a much smaller latent space (50 by default).

The encoder latent representation ***z*** is decoded with a 2 layers linear decoder network *D* : ℝ^*l*^ → ℝ^*g*^ to create a reconstruction of the original spliced and unspliced inputs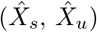 learning a function that maps from latent space to gene space.

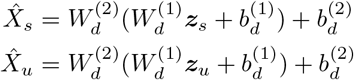

### Pseudotime encoding

The output of the first layer of the encoder is also used to estimate the pseudotime *t* of the cells, such that:

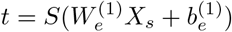

Where *S*(*x*) is the sigmoid function. Both pseudotime and latent representation of the cell are used to train nonlinear neural ODEs.

### Neural ODE

Once we project the cells into a lower-dimensional space, we use a set of neural ODEs to describe the cellular dynamics in this latent space

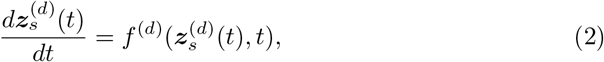

where *f*^(*d*)^ : ℝ^*l*^→ ℝ^*l*^ is a two layers neural network that takes *z*_*s*_ ∈ ℝ^*l*^ (with l = 50 by default) as input. The network uses the Exponential Linear Unit (ELU) function as the nonlinear activation function, and has a hidden layer dimension of 100. This neural ODE describes the dynamics of cells from particular sample *d*. Using the nonlinear ODEs in the latent space helps to capture the essential and complex dynamics of cells in this space.

Notice that, since we are learning both the pseudotime and the generator function of the dynamics (the *f* function), one could reverse simultaneously the pseudotime and the sign of *f* without altering the trajectories. This ambiguity may be resolved either by prior knowledge or by inverting both signs *a posteriori*.

### Linearisation and interpretation

For any dynamical process governed by a nonlinear ODE, we can understand the local behavior of the system by linearizing the ODE around a given point. In this case we choose as the point about which to linearise the average of the encoded state of (a group of) cells; the linearized system becomes:

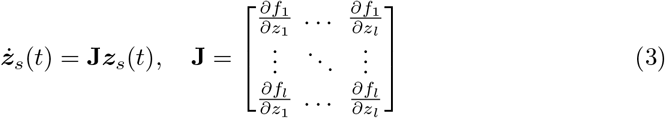

where the matrix of partial derivatives **J** is the Jacobian matrix. We perform *eigenanalysis* on the Jacobian matrix. The eigenvalues give the important directions of the correspondent eigenvectors in the latent space, then we use the decoder function to map these important directions in the latent space to the gene space. The output of this decoder is a list of ranked genes based on their importance in a particular direction.

The main issues related to using neural networks are interpretability and robustness, the former is not a problem because NeuroVelo uses both linear embedding and projection, thus the interpretation of the latent space into the gene space is straight-forward. The robustness part is more of a concern because a different initialization leads to a different local minima which makes biological interpretation uncertain. Here we propose a geometrical and statistical approach that gives a solid gene list by using many trained models.

The first step is to find the set of eigenvectors from other models that align the most with an eigenvector of a current model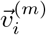, the alignment is based on the absolute value of the cosine similarity. This set of the most aligned eigenvectors is given by the following equation:

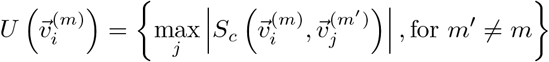

where *S*_*c*_ is the cosine similarity function and *m*^*′*^ is the index of the trained model.

We take the ranked genes of the aligned eigenvectors, and we average the ranks gene-wise. The output is a list of genes ranked based on the average rank across the aligned eigenvectors from different trained models. This indicates that the genes that are at the top of the list hold significant importance for the dynamics learned by various models and vice versa. A set of different analyses can be performed on these average-ranked genes for validation.

### GRN construction

As shown in equation 3, the non-linear latent dynamics are approximated by the linearized system ***ż***_*s*_(*t*) = **J*z***_*s*_(*t*).

This **J**, is obtained by linearizing the latent dynamical system around an average of group of cells, this could be all the cells -as done for benchmarking against other GRN methods- or by cell-type or cell pseudotime ordering -as done for the time dependent GRN in supplementary material-. By mapping the latent representation to the gene space, we can show that this system can be written as:

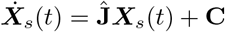

where **C** is a bias term that does not depend on X. We call **Ĵ** = **W**_**d**_**JW**_**e**_ the decoded Jacobian, **W**_**d**_ and **W**_**e**_ are the weight of the decoder and encoder networks respectively.

The matrix **Ĵ** ∈ ℝ^*g×g*^ (where *g* is the number of genes) captures the effective interactions between gene expression levels-i.e., the element Ĵ_*ij*_ tell us how a change in the expression of gene *j* influences the rate of change of gene *i*. A positive Ĵ_*ij*_ suggests that gene *j* activates gene *i*, while a negative value suggests that gene *j* represses gene *i* To construct a graph from the matrix **Ĵ** we take a directed graph *G* = (*V, E*) approach, such that each node *v*_*i*_ correspond to a gene *i* and a directed edge (*v*_*j*_ → *v*_*i*_) is weighted based on the magnitude of Ĵ_*ij*_.

### GRN evaluation: ChIP-seq data processing

The main challenge with benchmarking GRN is to establish the ground truth network to compare it with, and taking into consideration that gene regulatory network depends on the cell-type/cell-line. We create a *gold standard* gene regulatory network using chip-seq data from ChIP-Atlas [25], and we take the MACS2 score as a signal of the transcription factor binding to a target gene. We look at potential genes where the transcription factor binds at ± 10*k* distance from TSS of that gene.

To compare the inferred GRN by the 3 methods with the gold standard one, we take a sub-graph containing only TFs-Target genes pairs that exist in the our ground truth and we compute the area under precision recall curve (AUPRC). We use different cell lines and different threshold on the MACS2 score to evaluate the AUPRC, and we report AUPRC ratio, which is AUPRC/Baseline(Random Guess).

For the case of NeuroVelo and because we are taking MACS2 score as ground truth, we cannot benchmark for the type of interaction -whether it is activation or repression-, thus we take the magnitude as weighted graph edge.

### Model training

The tasks learnt by neural networks are based on what loss function the network is optimizing. To ensure that the cellular dynamics learned by the network have biological meaning, we propose a loss function that puts biophysical constraints on the data and does not require any parameter fine-tuning.

The first part of the loss is the usual mean squared error (MSE) of autoencoders network, we want our spliced and unspliced inputs to match their reconstructed counterparts from the decoder and so the first loss part is a sum of MSE of spliced and unspliced reads.

The second part of the loss is what puts a biophysical constraint on the neural ODEs in the same fashion of physics-informed neural networks. The idea is to make the derivative of latent spliced reads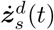 align with the splicing dynamics from the RNA velocity but in latent representation (*e*^*β*^***z***_*u*_ − *e*^*γ*^***z***_*s*_), There is an exponential of β (and γ) because they are analogy of splicing and degradation rate and they must be positive.

The final loss is written as follows:

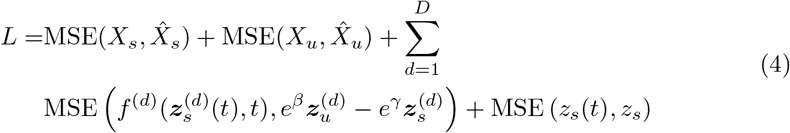

Where the sum over *d* is the sum over specific group of cells or treatments. 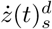is the function that describes the splicing dynamics of a sample *d* in the latent space. 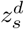and 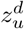 are the projected spliced and unspliced reads of the sample *d*. Notice that the third term in the loss enforces the RNA-velocity constraint, so that the role of the transcription rate is modelled flexibly and learnt directly from the data. The last term in the loss aim to align the latent dynamics with the latent representation, due to the linearity of the model, this is equivalent to MSE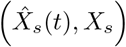 where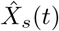 is the decoded latent dynamics *z*_*s*_(*t*).

In this way, we are able to investigate complex and specific cellular dynamics by learning splicing dynamics constrained nonlinear ODEs in a reduced space and without the need to fine tune parameters on the level of the loss function as well.

### Parameters selection

RNA velocity estimation can be very sensitive to the processing done such as constructing kNN graph as shown in [14] to use “moments” instead of raw reads. In addition, using neural network to estimate RNA velocity increases the number of parameter combinations, such as learning rate, optimizer, batch size and network size (dimensions). Moreover, for the same set of parameters, different initializations can be stuck in different local minima, leading to convergence in any case, but to different velocity estimates.

In address this issue, we propose a metric, *Velocity Alignment* which uses the same cosine similarity measure as *ICCoh* but is applied to velocities estimated using models trained with different random seeds.

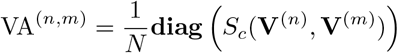

*V* ^(*n*)^ and *V* ^(*m*)^ are RNA velocity estimated from different trained model (*n*) and (*m*) and N is the total number of cells. The **diag** takes the diagonal elements of the cosine similarity measure, meaning that we are measuring how the same cell velocity from 2 different models is aligned.

The idea is to select the set of parameters with the highest cosine similarity. This approach ensures that even if the model converges to a local minimum, the velocity estimated from different seeds is aligned.

It is important to point that this metric should be computed for models with different random initializations but for the same set of other parameters.

### Implementation

NeuroVelo is implemented in Python machine learning framework `PyTorch 2.0` and using `torchdiffeq` package for ODE solvers in PyTorch.

Velocity field visualizations are done with `scvelo` except for the mouse lung cancer dataset it was done using `PhyloVelo` package. Gene networks visualizations are done with `networkx`

Gene set enrichment analysis is done with *“GSEApy: Gene Set Enrichment Analysis in Python”* `gseapy`. Passing a dataframe of genes and their average rank as a ranking metric to `prerank` function. The enrichment analysis plots are all generated by the same package.

The metrics *ICCoh* and *CBDir* are borrowed from [12]

### Datasets

#### Mouse erythroid

The human bone marrow dataset is downloaded from ScVelo package `scv.datasets.gastrulation_erythroid()`. The dataset is already processed and contains spliced and unspliced counts. We picked the top 2000 highly variables gene and we used counts training.It takes 524 seconds, 100 epoch for training and using CPU model: Intel(R) Xeon(R) Gold 6238R CPU @ 2.20GHz.

#### Mouse lung cancer

This dataset is CRISPR/Cas9-based lineage tracing dataset from a genetically-engineered mouse model (GEMM) of lung adenocarcinoma. We used it to compare NeuroVelo to PhyloVelo, mainly using two tumors (3726 NT T1 and 3435 NT T1).

We requested the spliced/unspliced reads from [20]. We picked the top 2000 highly variables gene and we used moments instead of counts for training.

#### Human bone marrow hematopoiesis

The human bone marrow dataset is downloaded from ScVelo package `scv.datasets.bonemarrow()`. The bone marrow is the primary site of new blood cell production or haematopoiesis. It is composed of hematopoietic cells, marrow adipose tissue, and supportive stromal cells. The dataset is already processed and contains spliced and unspliced counts. We picked the top 2000 highly variables gene and we used moments instead of counts for training.It takes 451 seconds, 100 epoch for training and using CPU model: Intel(R) Xeon(R) Gold 6238R CPU @ 2.20GHz

#### Pancreatic endocrinogenesis

Pancreatic endocrinogenesis dataset is downloaded from ScVelo package `scv.datasets.pancreas()`. Endocrine cells originate from endocrine progenitors found within the pancreatic epithelium. These progenitors differentiate into four primary cell types: α-cells that produce glucagon, β-cells that produce insulin, δ-cells that produce somatostatin, and ε-cells that produce ghrelin. The dataset is already processed and contains spliced and unspliced counts. We picked the top 2000 highly variables gene and we used counts training.It takes 185 seconds, 100 epoch for training and using CPU model: Intel(R) Xeon(R) Gold 6238R CPU @ 2.20GHz

## Code availability

The source code for NeuroVelo is available on GitHub: https://github.com/idriskb/NeuroVelo. Along with tutorial notebooks on how to run the method for different datasets, posts analysis for extracting the most important genes and gene regulatory network construction.

## Supplementary Figures

**Fig. S1.**
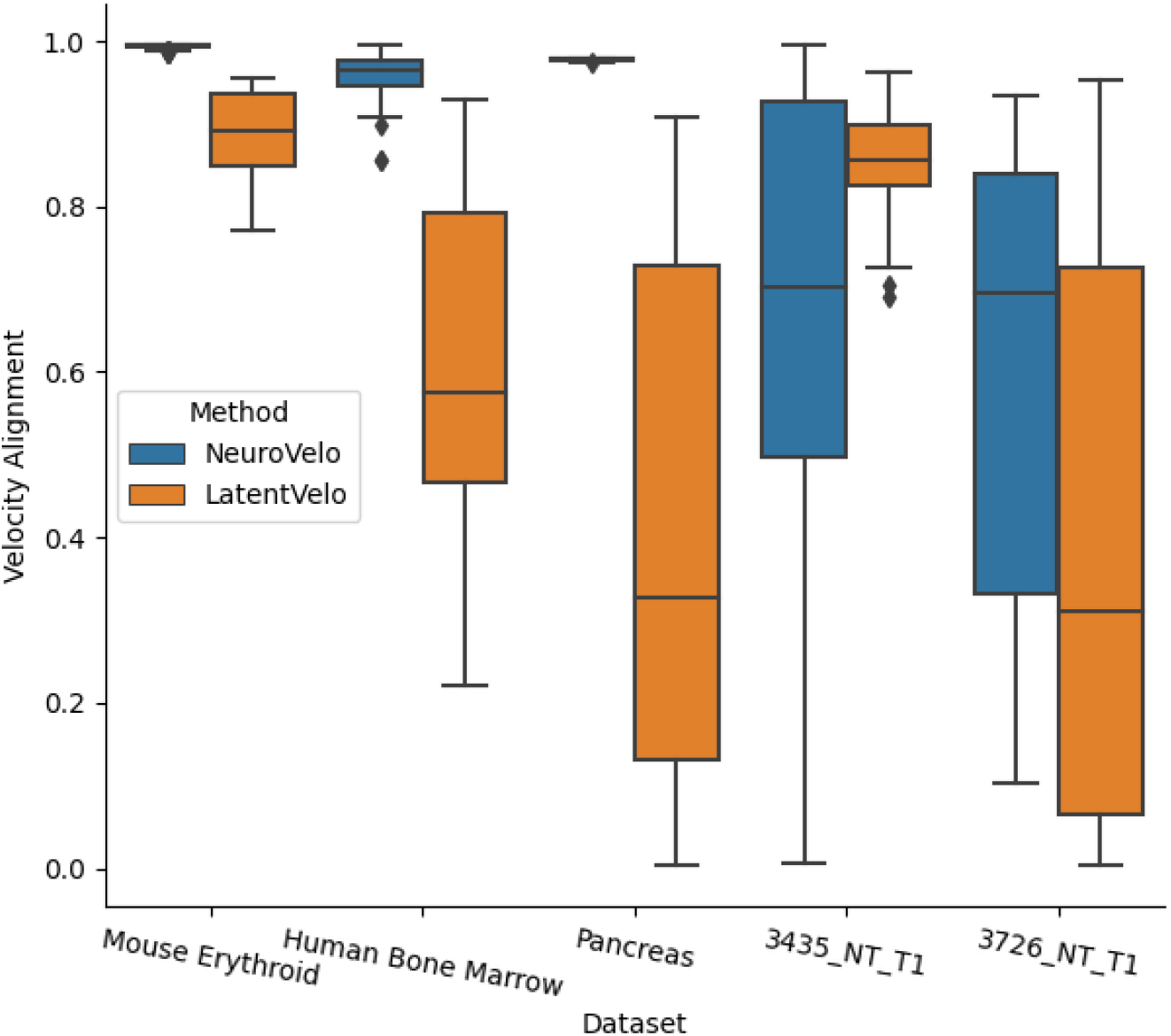
Comparison on velocity alignment, each point is the velocity alignment of a pair of trained models.

**Fig. S2.**
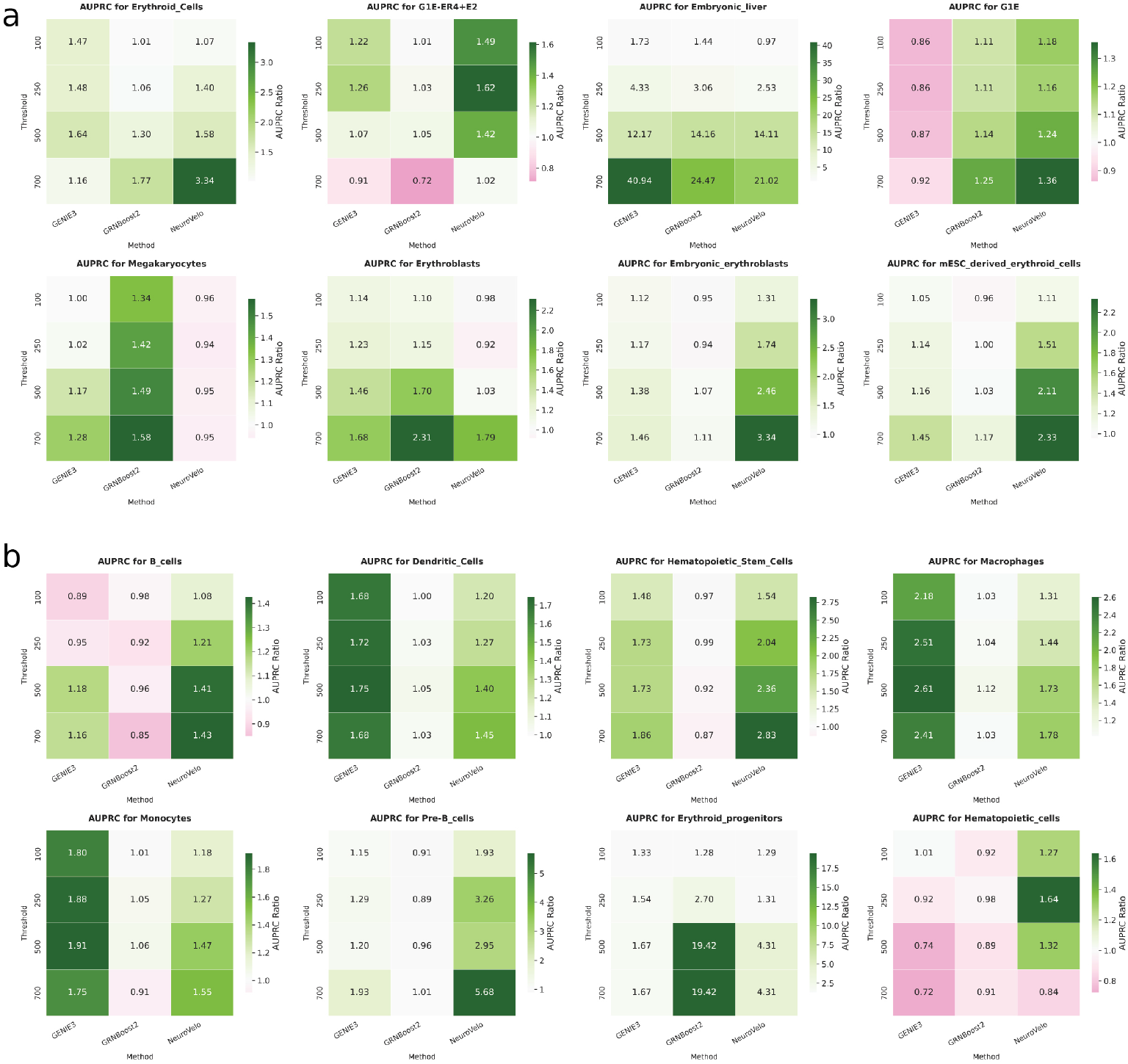
Comparison on AUPRC ratio for GRN inference performance at different threshold of MACS2 score. (a) Mouse erythroid dataset, (b) Human bonemarrow dataset

**Fig. S3.**
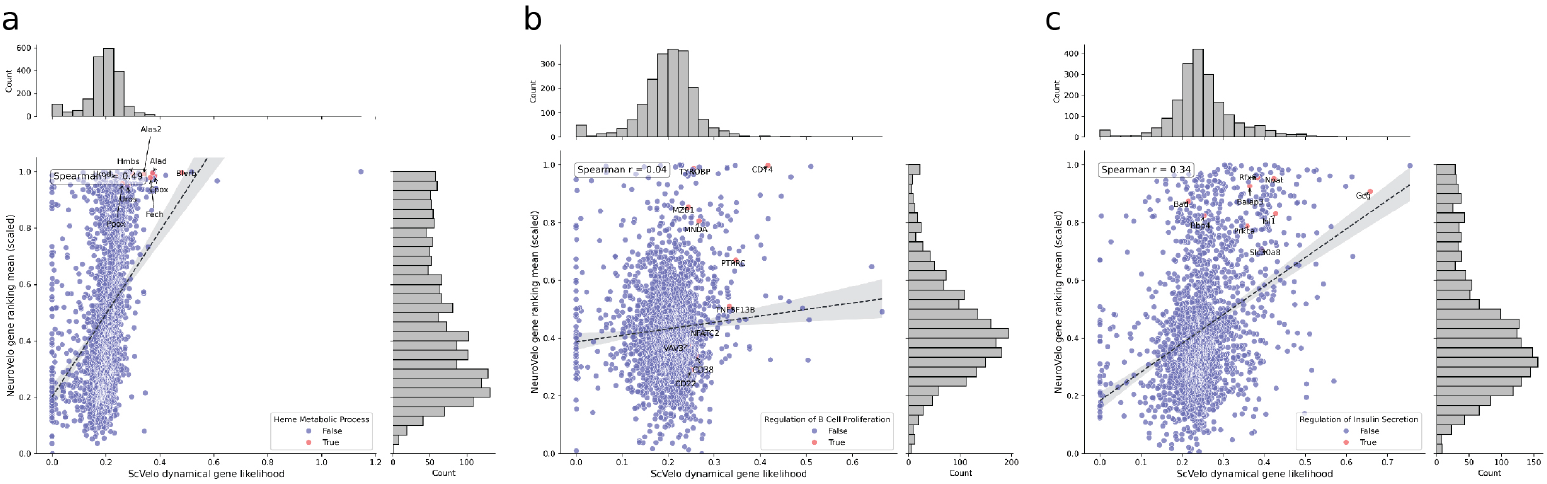
Comparison on spearman correlation for genes driving dynamics from ScVelo and NeuroVelo on (a) Mouse erythroid, (b) Human bone marrow, (c) Pancreas datasets.

## Supplementary Tables

**Table ST 1:**
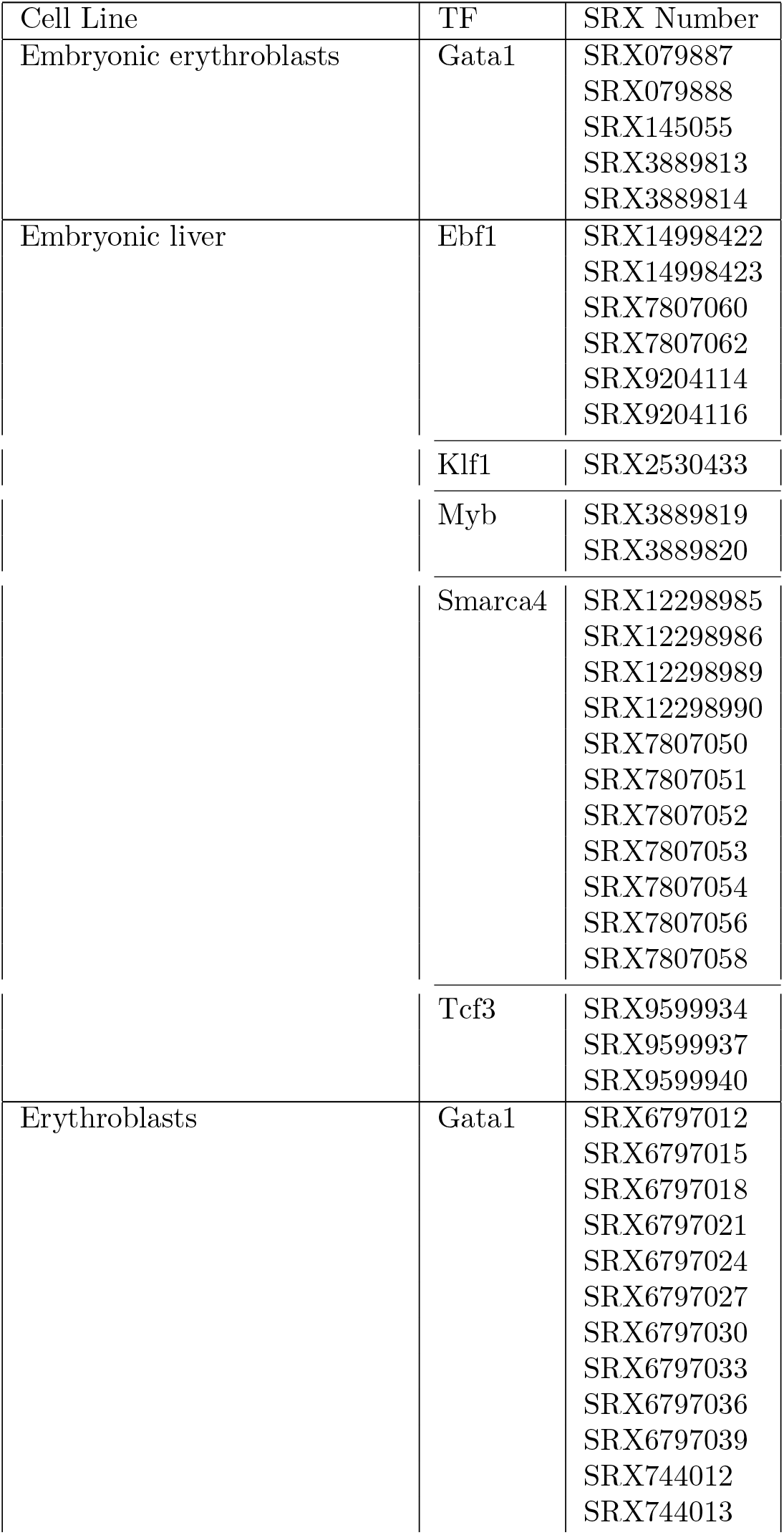
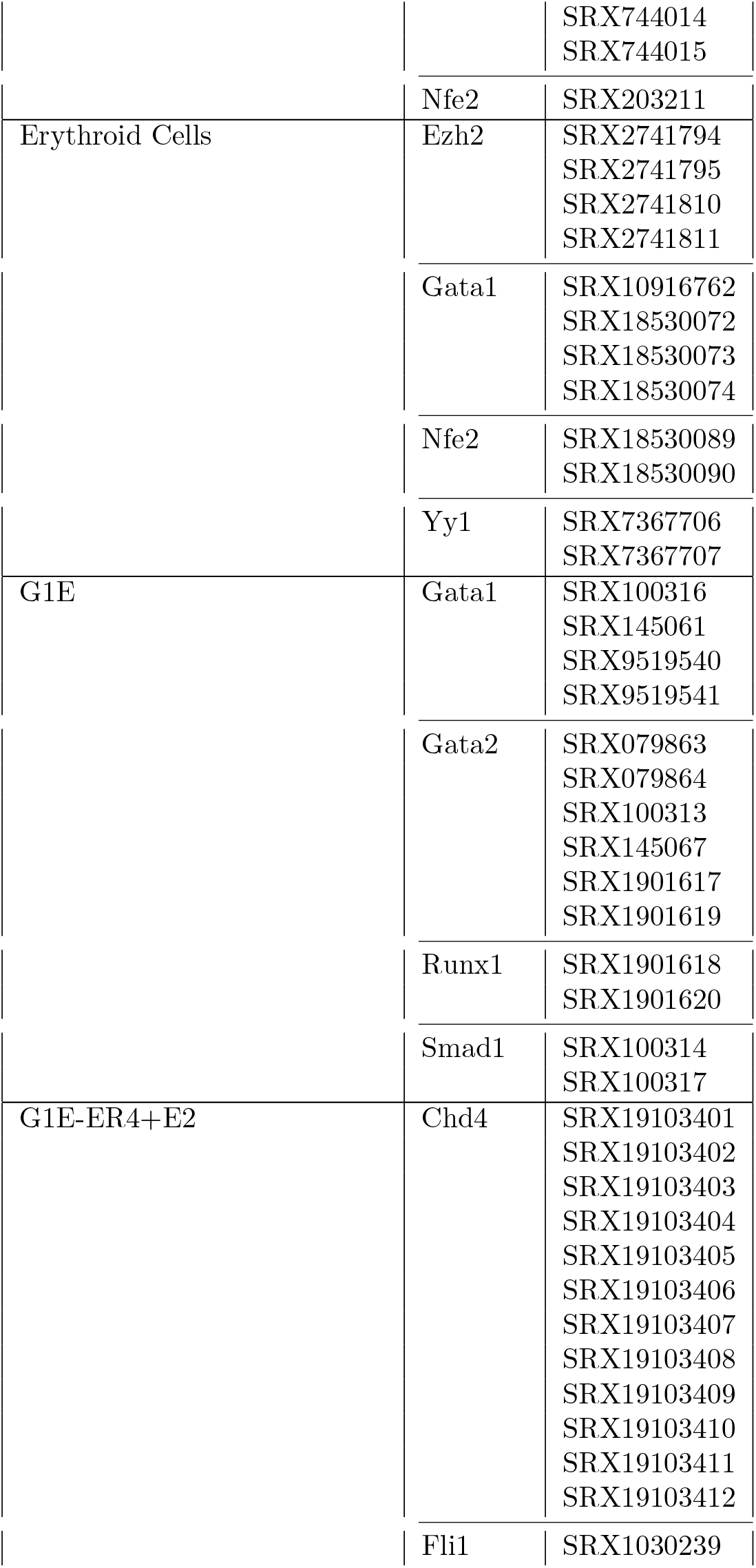
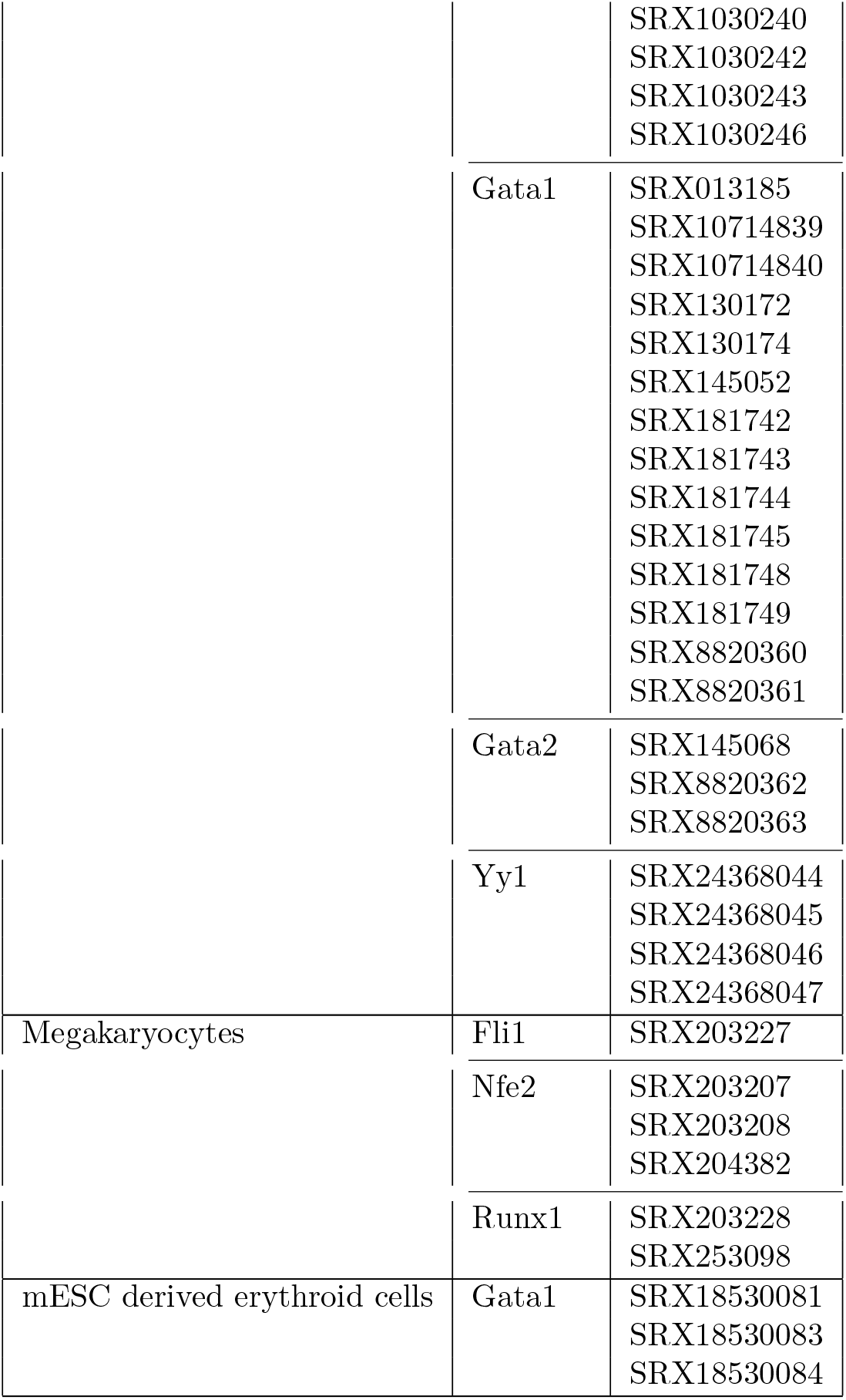
Summary of transcription factors (TFs) and their associated SRX accession numbers across different cell lines for mouse erythroid dataset.

**Table ST 2:**
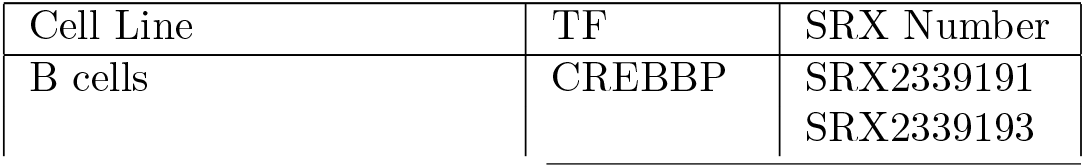
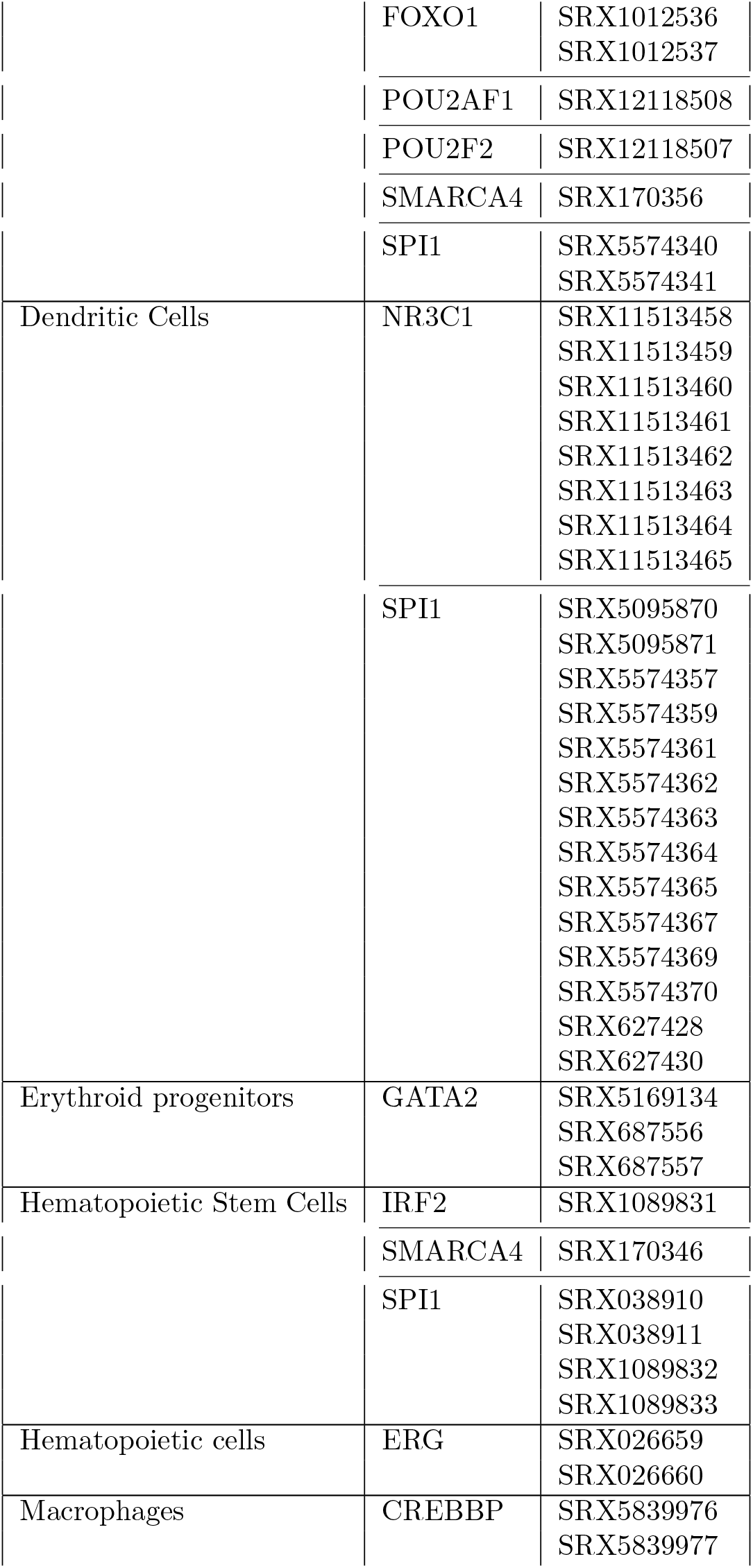
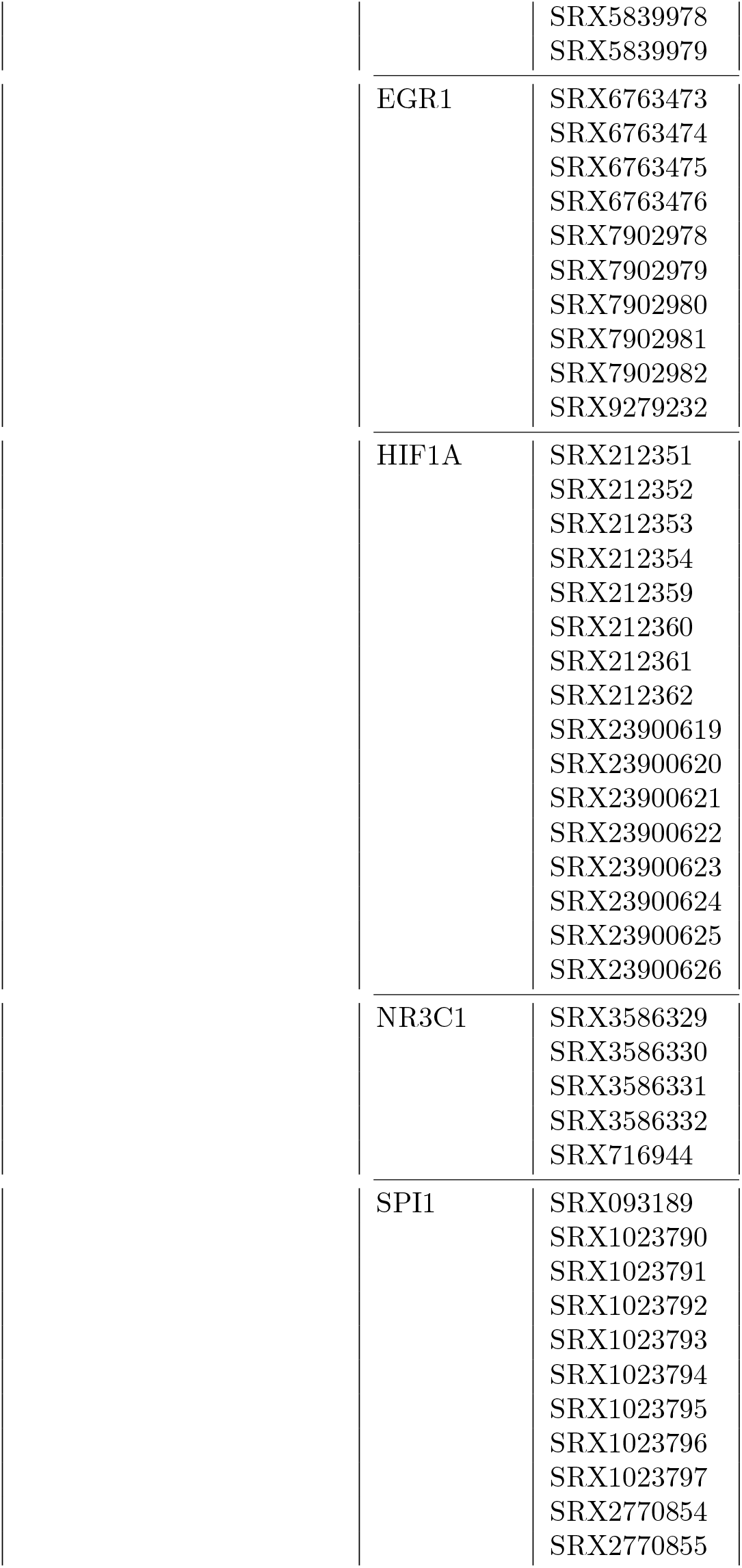
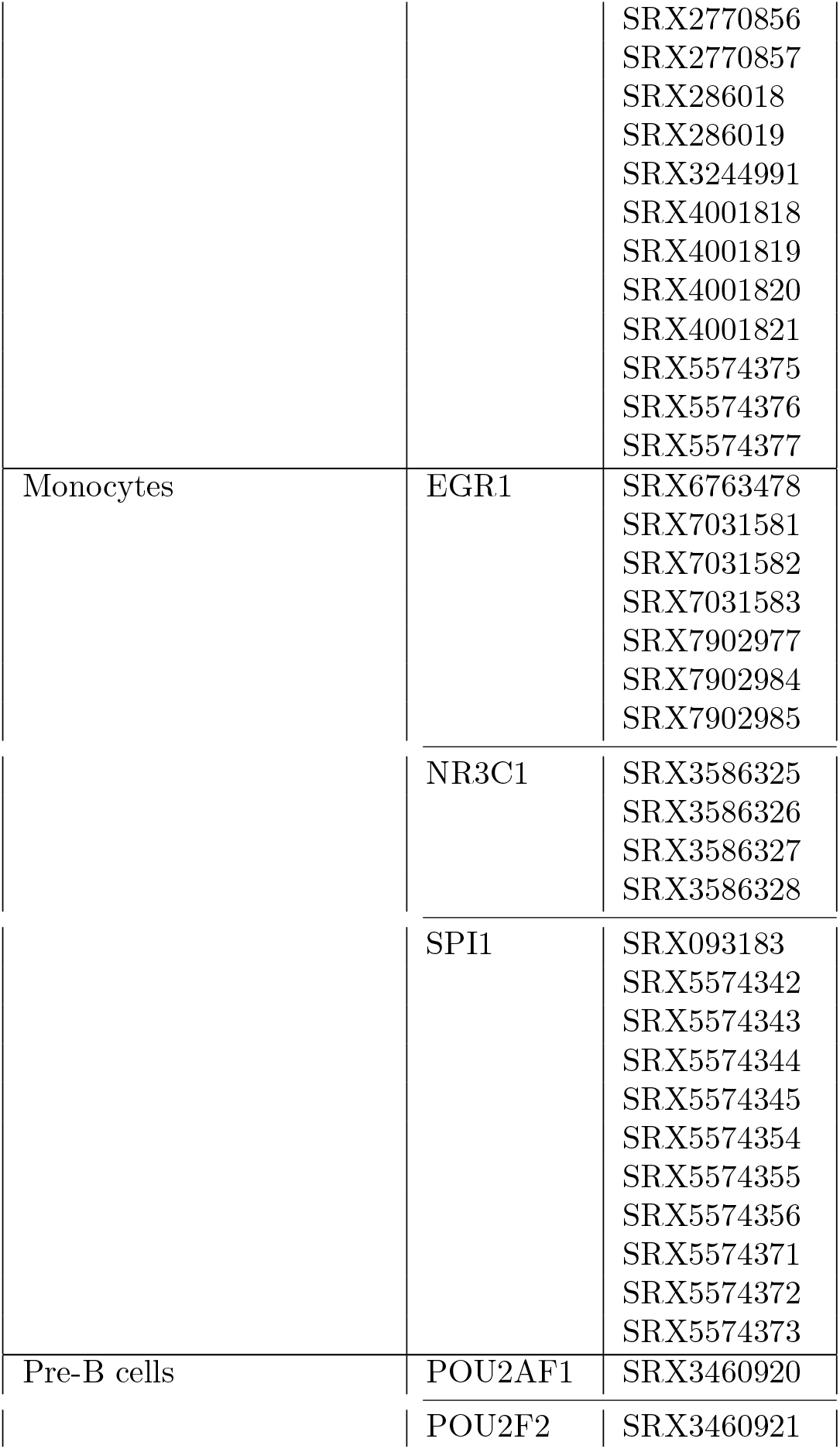
Summary of transcription factors (TFs) and their associated SRX accession numbers across different cell lines for human bone marrow dataset.

